# A High-efficacy CRISPRi System for Gene Function Discovery in *Zymomonas mobilis*

**DOI:** 10.1101/2020.07.06.190827

**Authors:** Amy B. Banta, Amy L. Enright, Cheta Siletti, Jason M. Peters

## Abstract

*Zymomonas mobilis* is a promising biofuel producer due to its high alcohol tolerance and streamlined metabolism that efficiently converts sugar to ethanol. *Z. mobilis* genes are poorly characterized relative to model bacteria, hampering our ability to rationally engineer the genome with pathways capable of converting sugars from plant hydrolysates into valuable biofuels and bioproducts. Many of the unique properties that make *Z. mobilis* an attractive biofuel producer are controlled by essential genes; however, these genes cannot be manipulated using traditional genetic approaches (e.g., deletion or transposon insertion) because they are required for viability. CRISPR interference (CRISPRi) is a programmable gene knockdown system that can precisely control the timing and extent of gene repression, thus enabling targeting of essential genes. Here, we establish a stable, high-efficacy CRISPRi system in *Z. mobilis* that is capable of perturbing all genes—including essentials. We show that *Z. mobilis* CRISPRi causes either strong knockdowns (>100-fold) using single guide RNA (sgRNA) spacers that perfectly match target genes, or partial knockdowns using spacers with mismatches. We demonstrate the efficacy of *Z. mobilis* CRISPRi by targeting essential genes that are universally conserved in bacteria, key to the efficient metabolism of *Z. mobilis*, or underlie alcohol tolerance. Our *Z. mobilis* CRISPRi system will enable comprehensive gene function discovery, opening a path to rational design of biofuel production strains with improved yields.

**IMPORTANCE:** Biofuels produced by microbial fermentation of plant feedstocks provide renewable and sustainable energy sources that have the potential to mitigate climate change and improve energy security. Engineered strains of the bacterium *Z. mobilis* can convert sugars extracted from plant feedstocks into next generation biofuels such as isobutanol; however, conversion by these strains remains inefficient due to key gaps in our knowledge about genes involved in metabolism and stress responses such as alcohol tolerance. Here, we develop CRISPRi as a tool to characterize gene function in *Z. mobilis*. We identify genes that are essential for growth, required to ferment sugar to ethanol, and involved in resistance to alcohol. Our *Z. mobilis* CRISPRi system makes it straightforward to define gene function and can be applied to improve strain engineering and increase biofuel yields.

---

***Z***. *mobilis* is a Gram-negative α-Proteobacterium with superlative properties for biofuel production (1–5), but poorly characterized gene functions (6). *Z. mobilis* is an efficient, natural ethanologen (7, 8) capable of fermenting glucose to ethanol at 97% of the theoretical yield (7, 9) with little energy spent on biomass (8, 10). Further, *Z. mobilis* is highly resistant to ethanol, up to 16% v/v (8). However, our ability to engineer strains of *Z. mobilis* that produce high yields of advanced biofuels, such as isobutanol (IBA), has been hindered by the lack of functional information for two key gene sets: metabolic and stress response/resistance genes. Genetic analysis to identify and characterize metabolic and stress response genes could allow us to engineer strains with increased flux toward IBA and away from ethanol (3, 11), as well as strains that are resistant to hydrolysate inhibitors such as acetic acid and various phenolic compounds (4).

*Z. mobilis* has a minimalistic metabolism with little functional redundancy (12–14). *Z. mobilis* converts sugars to pyruvate *via* the Entner-Doudoroff (ED) pathway, rather than the more commonly used but less thermodynamically favorable Embden–Meyerhof–Parnas pathway (EMP; (15–18)). Metabolic models based on *Z. mobilis* ZM4 genome sequences (12–14, 18) revealed that central metabolic pathways such as EMP glycolysis and the tricarboxylic acid (TCA) cycle are missing key enzymes (e.g., 6-phosphofructokinase and 2-oxoglutarate dehydrogenase, respectively), further limiting its metabolic plasticity. Because of its streamlined metabolism, many metabolic genes are predicted to be essential for growth in *Z. mobilis*.

Another defining feature of *Z. mobilis* physiology is the production of large quantities of hopanoids (19); *i.e.*, triterpenoid lipids that provide resistance to environmental stresses in bacteria (20–22). Hopanoids are thought to act by altering membrane fluidity and permeability, analogous to the action of cholesterol—also a triterpenoid lipid—on eukaryotic membranes (23–25). Although other bacteria make hopanoids, *Z. mobilis* produces them in higher quantities, with the number of hopanoids nearly matching that of phospholipids in the cell during peak production conditions (19). Hopanoids are thought to be essential to *Z. mobilis*, as chemical inhibition of enzymes involved in hopanoid precursor biosynthesis inhibits growth (26, 27) and transposon insertions into hopanoid biosynthesis genes result in cells with both a wild-type and transposon mutant allele (*i.e.*, *hpn*+/*hpn*::Tn strains (28, 29)). Consistent with having an essential role in *Z. mobilis* physiology, hopanoid production is correlated with ethanol content of the growth medium (30), and mutations in hopanoid biosynthesis genes increase sensitivity to ethanol (28). Whether hopanoids are required for resistance to additional stresses, such as hydrolysate toxins or alcohols other than ethanol is unknown.

CRISPRi—the use of programmable guide RNAs and Cas proteins (31, 32) or complexes (33, 34) lacking nuclease activity to repress transcription of target genes—is capable of probing the functions of essential genes. This is because CRISPRi knockdowns are inducible and titratable (31, 32, 35), separating the steps of strain construction and gene phenotyping. CRISPRi has been used to phenotype essential genes in multiple bacterial species (36), define chemical-gene interactions (35, 37), cell morphology phenotypes (35, 38, 39), host genes involved in phage life cycles (40), novel gene functions (35, 38), and essential gene network architecture (35), among others. To facilitate essential gene phenotyping by CRISPRi in diverse bacterial species, we recently developed “Mobile-CRISPRi”—a suite of modular CRISPRi vectors based on the extensively studied type II-A CRISPR system from *Streptococcus pyogenes* (*i.e.*, *Spy* dCas9; Fig. 1a) that are transferred by mating and site-specifically integrate into the genomes of recipient bacteria (41, 42). We demonstrated integration and knockdown in species ranging from Gram-negative γ-Proteobacteria to Gram-positive Firmicutes (41); although we did not test species from α-Proteobacteria. *Z. mobilis* contains a native type I-F CRISPR system that has been co-opted for efficient genome editing as well as CRISPRi in a Cas3 nuclease-deficient background (34); however, CRISPRi using the native system showed low knockdown efficacy (4-5 fold maximum) and was not inducible as constructed, limiting its usefulness in targeting essential genes. We reasoned that importing the heterologous *Spy* dCas9 system would result in stronger knockdowns, as is seen in other species (31, 35, 41).

**Fig 1.**
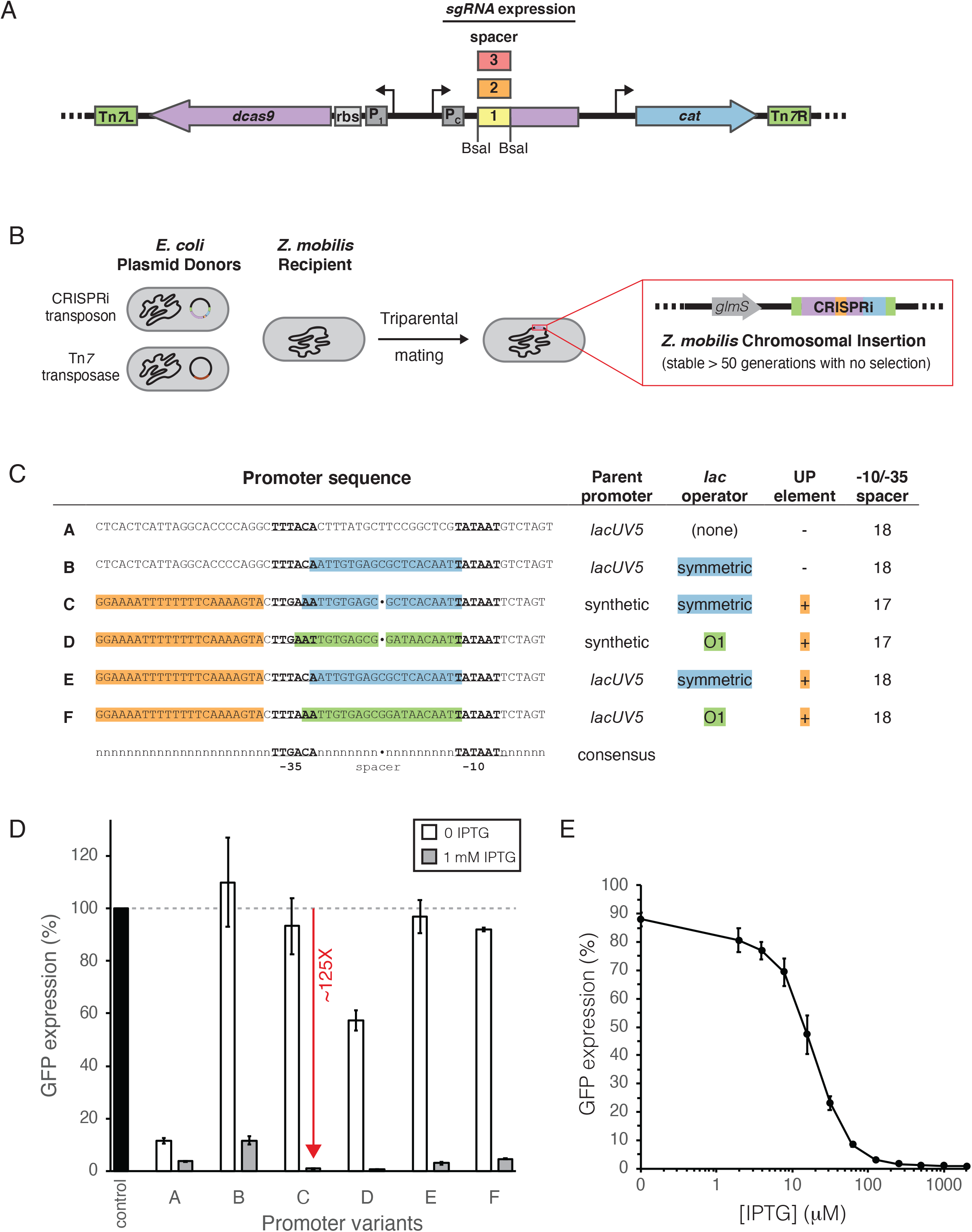
Mobile CRISPRi system for transcriptional repression optimized for *Zymomonas mobilis*. (A) Modular *Z. mobilis* CRISPRi system encodes dCas9, sgRNA, and antibiotic resistance cassettes on a Tn*7* transposon. The promoter (P_1_) and ribosome binding site (rbs) for dCas9 as well as the promoter (P_C_) for the sgRNA have been optimized for *Z. mobilis*. DNA encoding the 20 nt variable region of the sgRNA can be cloned (individually or libraries) in between the BsaI sites. (B) CRISPRi expressing strains are constructed by triparental mating of *E. coli* donor strains (one harboring the Mobile CRISPRi plasmid and another harboring a plasmid expressing the Tn*7* transposase) with *Z. mobilis*. The CRISPRi expression cassette will be stably incorporated onto the *Z. mobilis* chromosome at the Tn*7 att* site located downstream of *glmS*. (C) Optimization of sgRNA expression. Six promoter sequences (A-F) based on either lacUV5 or a synthetic promoter were incorporated into the CRISPRi system. Alignment is to the *E. coli* σ^70^ consensus promoter with the −10 and −35 core promoter elements underlined and shown in bold, the lac operator locations highlighted in green or cyan, and the UP element highlighted in yellow. (D) Comparison of *Z. mobilis* Mobile CRISPRi sgRNA promoter variants. A GFP expression cassette was cloned into the PmeI site and an sgRNA targeting GFP (or a non-targeting control) was cloned into the BsaI sites. Cultures were diluted 1:1000 and incubated in medium with 0 or 1 mM IPTG for ~10 doublings prior to measurement of GFP expression. Expression was normalized to a non GFP-expressing strain. Standard deviation between 4 biological replicates is shown. (E) Expression of *Z. mobilis* CRISPRi system is inducible over a range of IPTG concentrations.

Here, we establish a stable and efficacious CRISPRi system for *Z. mobilis* based on *Spy* dCas9. We demonstrate strong (>100-fold) or partial knockdown of gene expression by using sgRNA spacers that are complementary or mismatched to target genes, respectively. We use *Z. mobilis* CRISPRi to demonstrate the essentiality of genes involved in metabolism and hopanoid biosynthesis. Further, we show that reduced expression of specific hopanoid biosynthesis genes leads to IBA sensitivity. Our *Z. mobilis* CRISPRi system will enable rapid characterization of gene function, accelerating rational engineering of the genome for advanced biofuel production.

## RESULTS

### Optimization of CRISPRi for *Z. mobilis*

To establish *Spy* dCas9-based CRISPRi in *Z. mobilis*, we first attempted to deliver a previously described, Tn*7*-based Mobile-CRISPRi “test” vector containing the gene encoding monomeric Red Fluorescent Protein (mRFP) and an sgRNA targeting mRFP (41) to wild-type *Z. mobilis* strain ZM4 *via* conjugation (Fig. 1a and b). We failed to obtain transconjugants with wild-type but succeeded at integrating Mobile-CRISPRi into the genome of a restriction deficient derivative strain (Lal *et al*., *in preparation*), consistent with Mobile-CRISPRi vectors containing multiple predicted recognition sites for *Z. mobilis* restriction enzymes (43). Thus, we used the restriction-deficient strain in all subsequent experiments (Table 1). Mobile-CRISPRi inserted into the *Z. mobilis* genome downstream of *glmS* as expected (Fig. S1) and was stable over 50 generations of growth in rich medium without selection. We next used a fluorimeter to measure CRISPRi knockdown of mRFP at saturating concentrations of inducer (1 mM IPTG), finding poor knockdown (2.4-fold; Fig. S2), although our measurements were complicated by the weak fluorescence of mRFP in *Z. mobilis*.

**Table 1.**
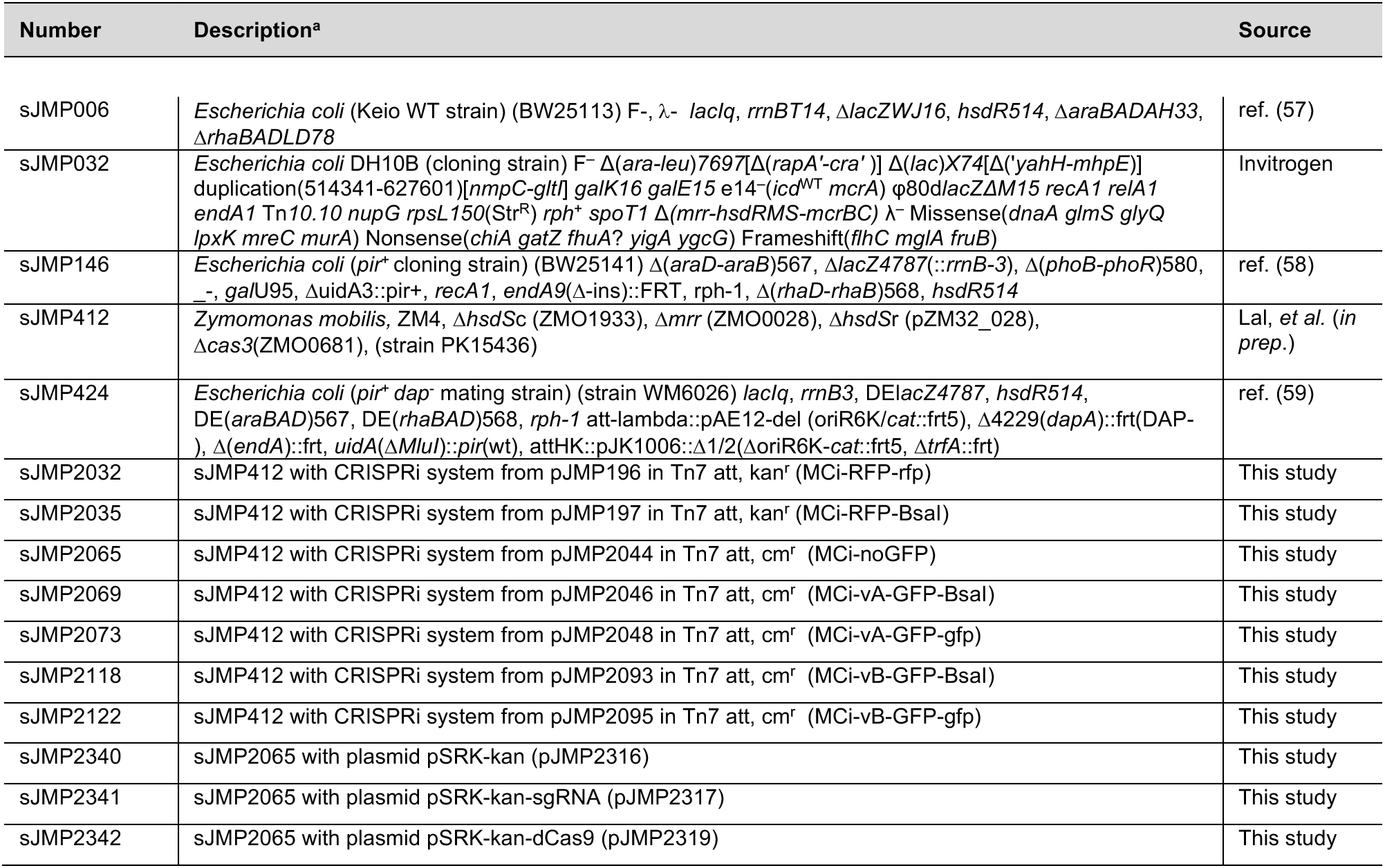
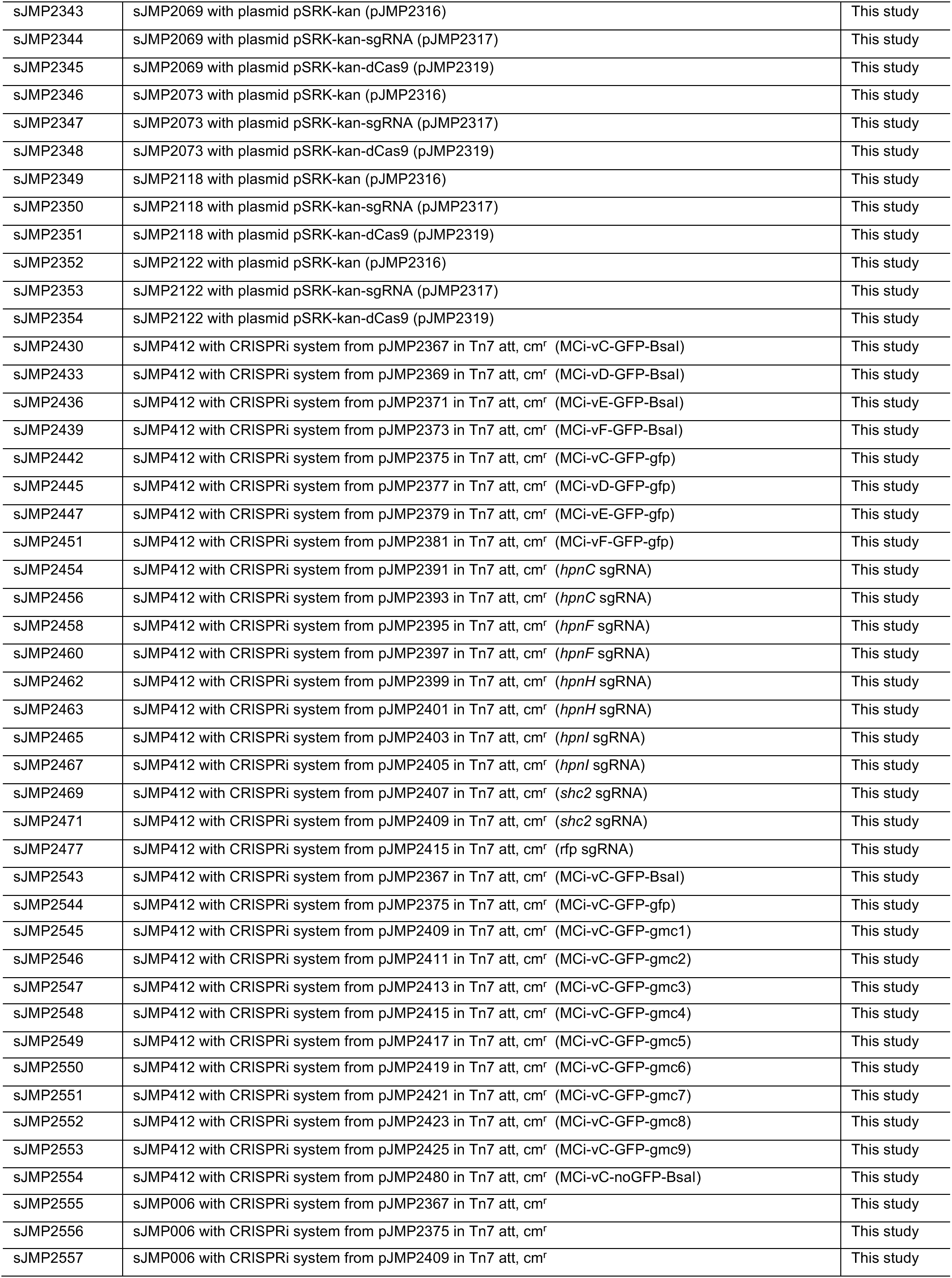
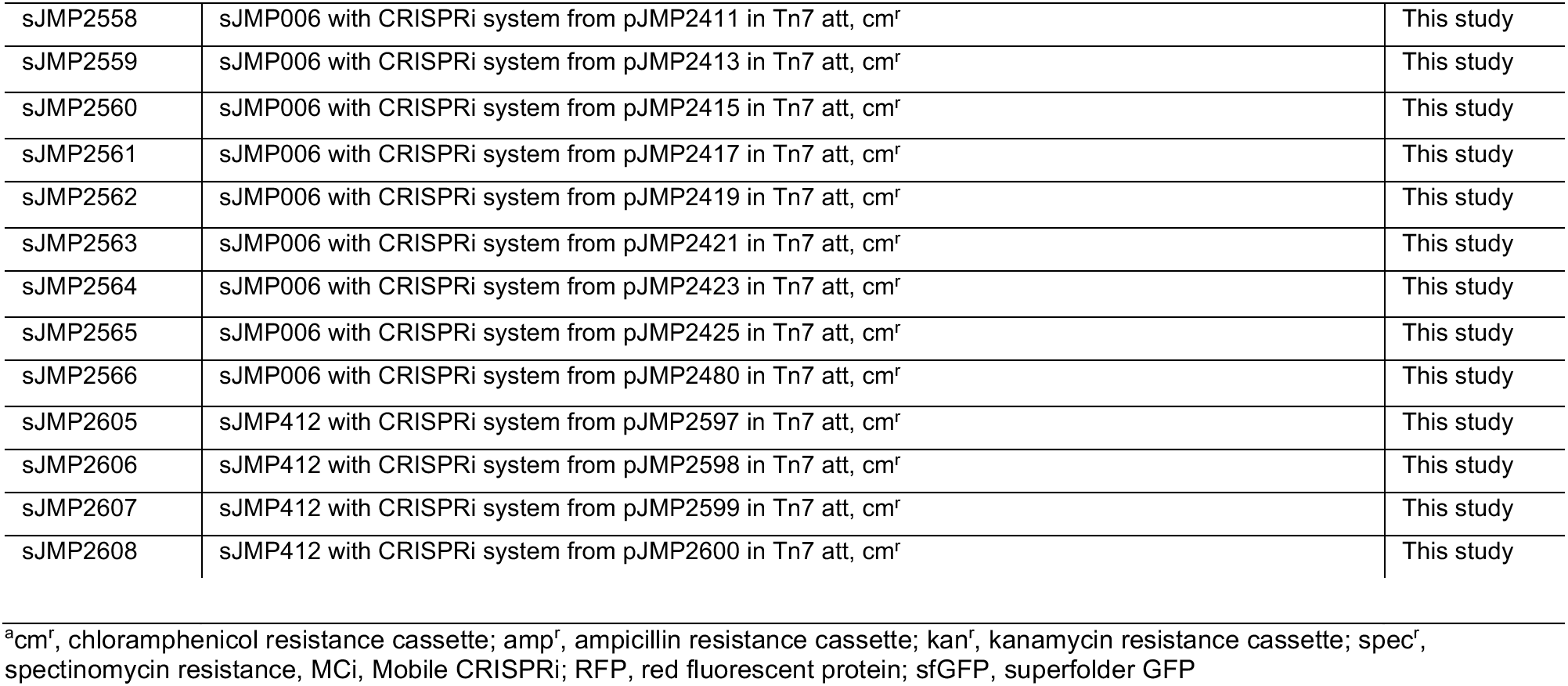
Strains.

To optimize CRISPRi function and improve knockdown detection, we took advantage of the modularity of Mobile-CRISPRi (Fig. 1a) to swap in biological “parts” that have been confirmed to function in *Z. mobilis* (44). We replaced mRFP with the gene encoding superfolder GFP (*sfGFP* (45)), expressed *dcas9* from a T7A1-derived promoter with a “strong” *Z. mobilis* ribosome binding site, and expressed an sgRNA targeting *sfGFP* from the *lacUV5* promoter (*i.e.*, promoter A; Fig. 1c). This CRISPRi system showed strong knockdown (28-fold) of sfGFP at saturating inducer, but also considerable leakiness (9-fold) without inducer (Fig. 1d, promoter A). Inserting a symmetric *lac* operator site (46) into the *lacUV5* promoter spacer (Fig. 1c, promoter B) resulted in no detectable leakiness but only modest knockdown (9-fold; Fig. 1d), suggesting that the concentration of either dCas9 or the sgRNA was limiting for knockdown. To determine the limiting factor, we expressed either the *sfGFP* sgRNA or *dcas9* from a multicopy plasmid in the context of CRISPRi with promoter B and found that sgRNA expression was primarily limiting knockdown (Fig. S3). Because *lacUV5* has the highest confirmed activity of any promoter measured in *Z. mobilis* (44) and because no consensus sequence exists in the literature for native *Z. mobilis* promoters, we built LacI-regulated synthetic promoters based on *Escherichia coli* σ^70^ consensus elements (47, 48) that we reasoned could increase sgRNA expression to a higher level than *lacUV5* (Fig. 1c, promoters C-F). All four synthetic promoters improved the knockdown properties of *Z. mobilis* CRISPRi, but promoter C—which features consensus UP and −10 elements with a near consensus −35 and an ideal spacer length (Fig. 1c)—provided the best combination of strong knockdown (125-fold) and negligible leakiness (~10-15%; Fig. 1d). Using CRISPRi with promoter C, we found that intermediate inducer concentrations enabled titration of knockdown activity (Fig. 1e). We conclude that Mobile-CRISPRi optimized for *Z. mobilis* is efficacious, inducible, and titratable.

### Mismatch-CRISPRi enables knockdown gradients in *Z. mobilis*

The relationship between fitness and gene expression varies by gene and is generally unknown (49, 50). This relationship is especially important to consider for essential genes, which have a fitness of zero at full knockdown but a large range of possible fitness values at intermediate levels of knockdown depending on the function of the gene product. Excessive knockdown of essential genes results in strains that grow poorly and are difficult to phenotype. Recent work from the Gross and Weissman labs has shown that systematically introducing mismatches between sgRNA spacers and target genes can generate knockdown gradients suitable for studying essential gene function (49, 50); we call this strategy “Mismatch-CRISPRi”. Mismatch-CRISPRi functions in mammalian cells (50) and diverse model bacteria (*i.e.*, *E. coli* and *B. subtilis* (49)), but has not been demonstrated in α-Proteobacteria. To test Mismatch-CRISPRi in *Z. mobilis*, we cloned the exact set of mismatched sgRNAs used by Hawkins and Silvis *et al*. to target *sfGFP* (49) into our *Z. mobilis* CRISPRi system with sgRNA promoter C (Fig 2a and b). Using these mismatched guides, we were able to generate a knockdown gradient of *sfGFP* that spanned nearly two orders of magnitude and contained multiple sgRNAs that caused intermediate knockdown levels at saturating inducer (Fig. 2c). Further, we introduced our *Z. mobilis* Mismatch-CRISPRi vectors into *E. coli* permitting a direct comparison of *sfGFP* knockdown in the two divergent species (Fig. S4). Consistent with a previous comparison between *E. coli* and *B. subtilis* (49), we found excellent agreement between *sfGFP* knockdown gradients in *E. coli* and *Z. mobilis* (R^2^ = 0.8). This demonstrates the broad utility of Mismatch-CRISPRi to predictably generate partial knockdowns and suggests that *Z. mobilis* CRISPRi may function well in multiple species.

**Fig 2.**
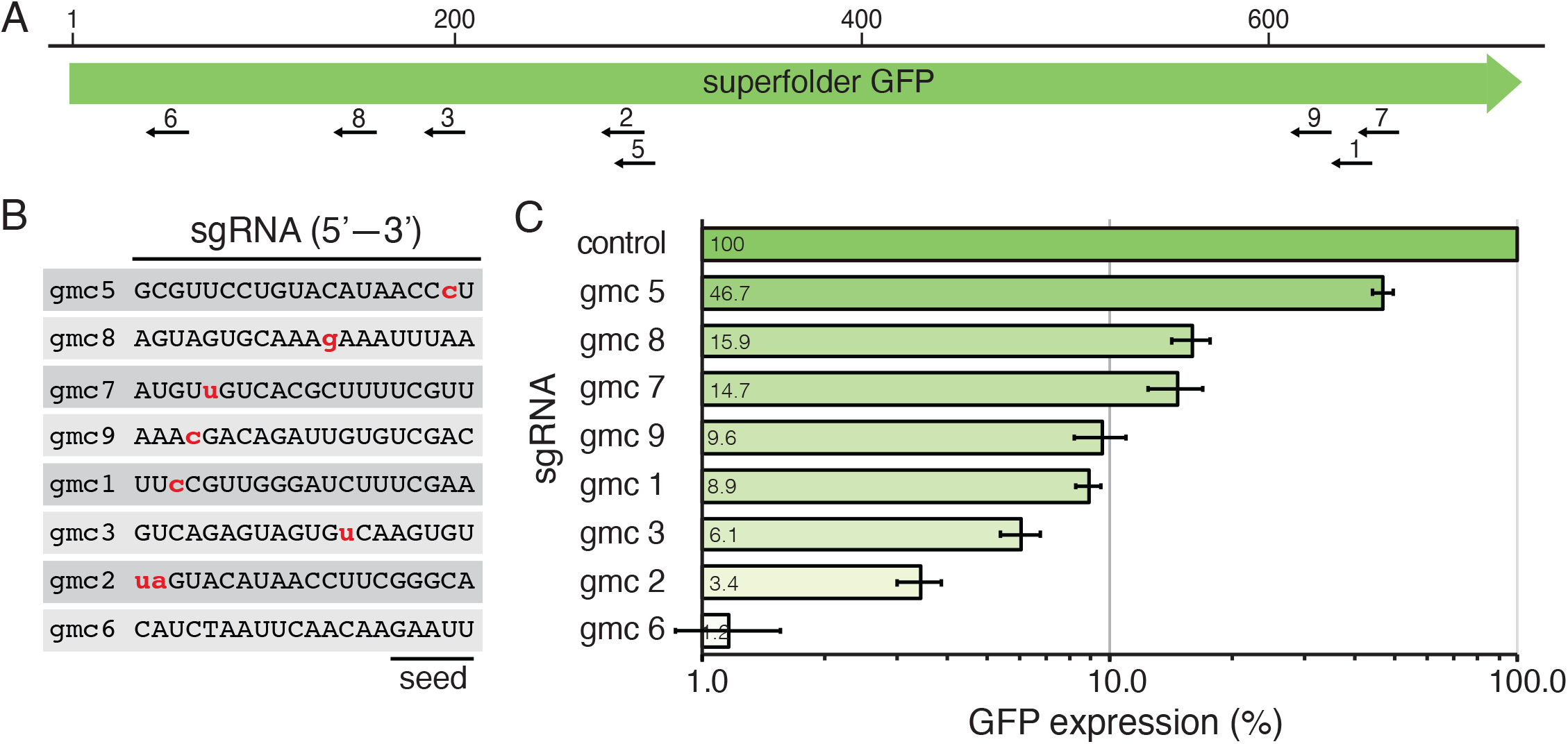
Variable repression using mismatch sgRNAs in the *Z. mobilis* Mobile CRISPRi system. (A) Location of sgRNA targets gmc1-9 on the GFP gene. Scale bar indicates nucleotides. (B) Sequences of the GFP-targeting sgRNAs with mismatches indicated in lowercase red. PAM proximal seed sequence is indicated. (C) Knockdown of GFP expression in *Z. mobilis* CRISPRi expression strains with mismatch sgRNAs. Control indicates a non-targeting sgRNA.

### *Z. mobilis* CRISPRi targets essential genes

To examine the efficacy of *Z. mobilis* CRISPRi in characterizing essential gene function, we first targeted *rplL* (ZMO0728)—an essential gene encoding the universally conserved ribosomal protein, L12 (51)—as a positive control. We found a greater than six orders of magnitude reduction in plating efficiency for strains expressing an *rplL* sgRNA versus a control strain expressing a non-targeting sgRNA at saturating inducer (Fig. 3a), indicating substantial loss of cell viability and relatively low levels of suppressor mutations that inactivate the CRISPRi system. Based on these results, we conclude that *Z. mobilis* CRISPRi is effective at assessing gene essentiality and allowing observation of essential gene knockdown phenotypes.

**Fig 3.**
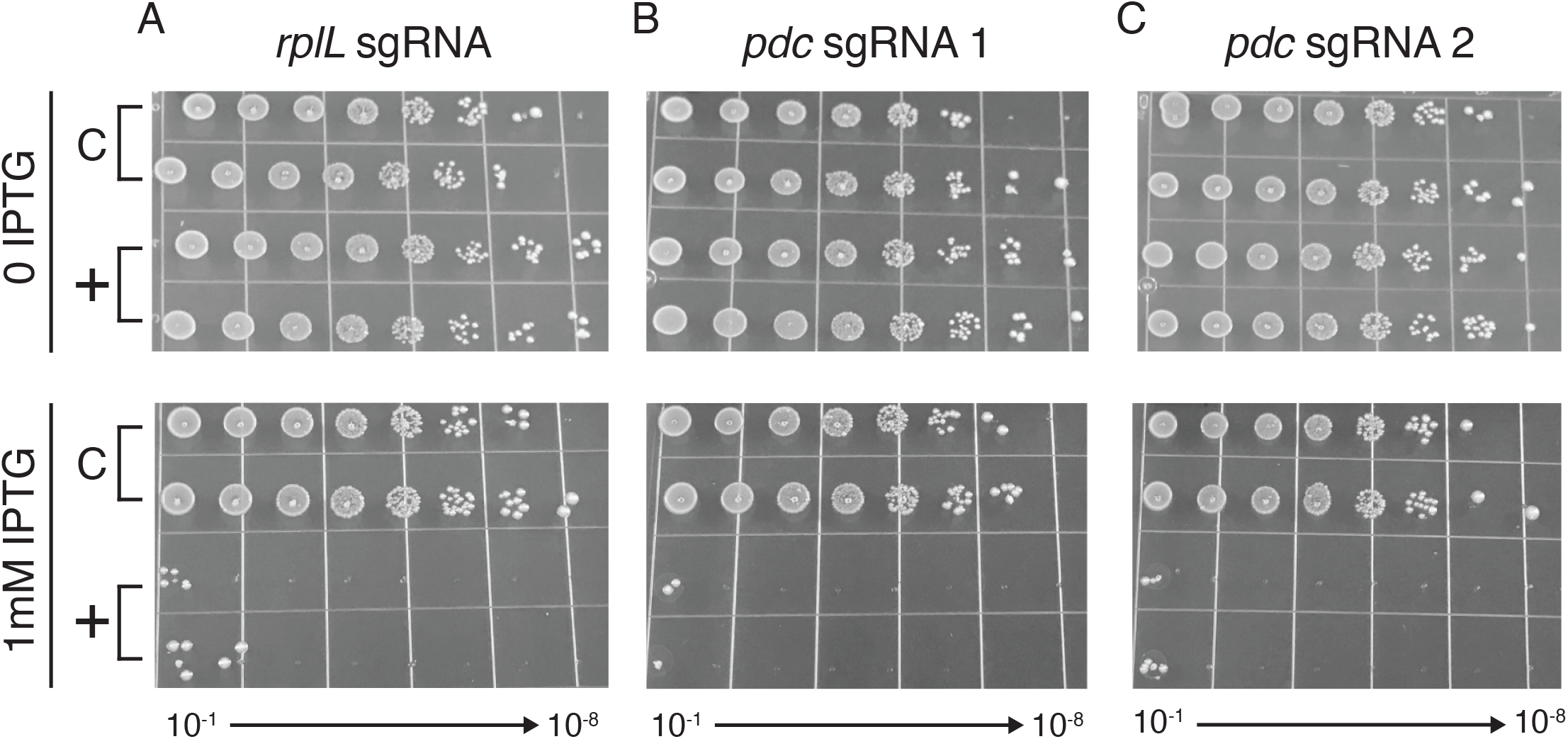
CRISPRi knockdown of endogenous essential genes. *Z. mobilis* strains with CRISPRi cassettes encoding sgRNAs targeting essential genes *rplL* (panel a) *and pdc* (panels b and c) were serially diluted 1:10 (10^−1^ through 10^−8^) and spotted on agar plates with either 0 or 1 mM IPTG. C indicates a non-targeting sgRNA.

Pyruvate decarboxylase, encoded by the *pdc* gene (ZMO1360), is a key metabolic enzyme in *Z. mobilis* that converts pyruvate into acetaldehyde—the penultimate step in ethanol production (52). Despite its important role in fermentation of sugars to ethanol, the *Z. mobilis* literature is conflicted about whether *pdc* is essential (5) or dispensable (53, 54) in aerobic conditions. To determine the essentiality of *pdc*, we used *Z. mobilis* CRISPRi with promoter C and an sgRNA targeting the 5′ end of the *pdc* coding sequence. We found a greater than six orders of magnitude loss in plating efficiency for the *pdc* knockdown strain at saturating inducer (Fig. 3b); this result was indistinguishable from the loss of fitness observed when we targeted *rplL*, suggesting that *pdc* is essential for aerobic growth. To confirm that our result was not due to off-target effects of CRISPRi, we tested a second, non-overlapping sgRNA targeting *pdc*, finding the same results (Fig. 3c). We conclude that *pdc* is essential for aerobic growth of *Z. mobilis*.

### Essentiality and IBA sensitivity phenotypes of hopanoid biosynthesis genes

Genes encoding hopanoid biosynthesis enzymes (*i.e.*, *hpn*/*shc* genes; Fig. 4a and S4) are thought to be essential in *Z. mobilis* based on growth cessation caused by small molecule inhibitors of Squalene-hopene cyclase (26) and the observation that strains with transposon insertions in *hpn* genes always also contain a wild-type copy of the gene (28). To further probe the essentiality of hopanoids, we targeted *hpn*/*shc* genes using *Z. mobilis* CRISPRi. Because CRISPRi blocks transcription of downstream genes in an operon (*i.e.*, polarity), we chose to target the first *hpn*/*shc* gene present in each operon (Fig. 4a, orange genes). We found considerable defects in plating efficiency for strains with sgRNAs targeting *hpnC* (ZMO0869), *hpnH* (ZMO0874), and *hpnI* (ZMO0972), consistent with a requirement of hopanoid synthesis for growth (Fig. 4b). In contrast, targeting the *hpnF* (a.k.a., *shc1*; ZMO0872) and *shc2* (ZMO1548) genes that both encode Squalene-hopene cyclase had no effect on plating efficiency, suggesting that they are functionally redundant under the conditions tested.

**Fig 4.**
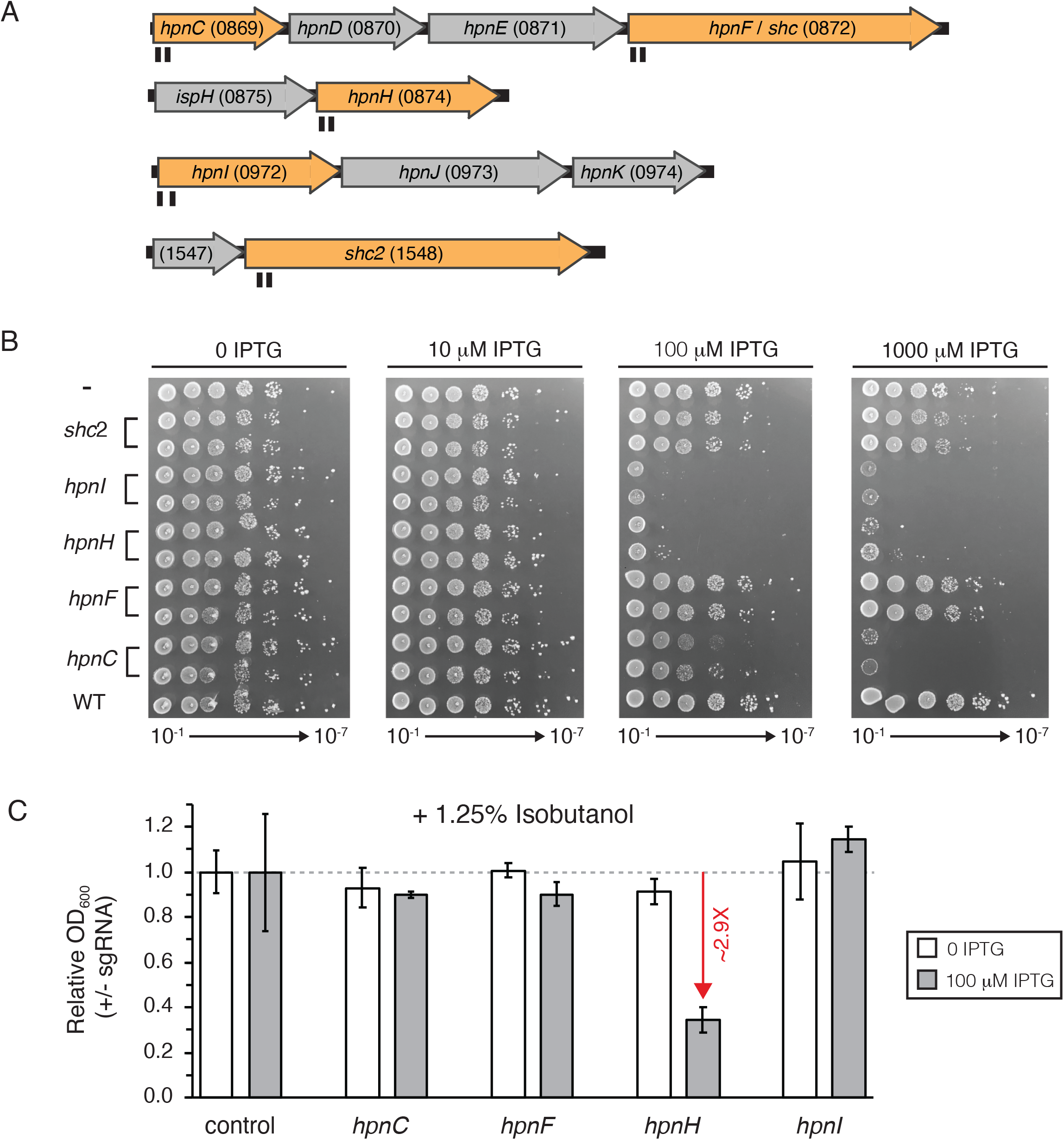
CRISPRi knockdown of hopanoid lipid synthesis related genes. (A) *Z. mobilis* strains were constructed with CRISPRi cassettes encoding sgRNAs targeting genes in hopanoid synthesis operons (and a non-targeting control). ZMOxxxx locus tag indicated after the gene name. Targeted genes (*hpnC*, *hpnF*, *hpnH*, *hpnI*, *shc2*) are shown in orange. Target positions of sgRNAs are shown in black under the genes. (B) Strains were serially diluted 1:10 (10^−2^ through 10^−8^) and spotted on agar plates with either 0, 0.1, or 1 mM IPTG. C indicates a non-targeting sgRNA. (C) Strains were diluted 1:1000 and grown ~10 doublings in liquid culture, aerobically in the presence of 1.25% isobutanol and 0 or 0.1 mM IPTG, prior to measurement of cell density (OD A_600_). Growth measurements were normalized to a strain expressing a non-targeting sgRNA. Standard deviation between 4 replicates is shown. Red arrow indicates fold change compared to control.

Classic studies of *Z. mobilis* physiology (27) and contemporary work using *hpn*+/*hpn*::Tn strains (28) have linked hopanoid production and ethanol concentration or resistance; however, it is unknown if hopanoids provide resistance to advanced biofuels, such as IBA. To examine the relationship between *hpn*/*shc* genes, we first determined the concentration of IBA needed to partially inhibit growth of *Z. mobilis* in sealed, deep 96 well plates. We found that addition of 1.25% v/v IBA to rich medium inhibited *Z. mobilis* growth by ~50% (Fig. S6). We then grew our CRISPRi strains targeting *hpn*/*shc* operons with a sub-saturating concentration of inducer (100 μM IPTG) for a limited number of generations in the presence or absence of IBA. Under these conditions, *hpnH* was the only knockdown tested that showed increased sensitivity to IBA, with a 2.9-fold reduction in final OD_600_ at the end of the growth period relative to a non-targeting control sgRNA strain. HpnH performs the first enzymatic step in hopanoid side chain synthesis (Fig. S5; (55)), suggesting that buildup of core hopanoids diploptene and diplopterol may compromise IBA tolerance in *Z. mobilis*. We conclude that hopanoid biosynthesis operons are essential in *Z. mobilis*, but knockdown of extended hopanoids does not alter IBA tolerance.

## DISCUSSION

The lack of genetic tools for bioenergy-relevant, non-model bacteria has slowed our progress toward engineering efficient production strains for advanced biofuels, such as IBA. Our optimized CRISPRi system for *Z. mobilis* overcomes this obstacle by enabling programmable, inducible, and titratable control over expression of both non-essential and essential genes. Because *Z. mobilis* CRISPRi also functions well in the distantly related bacterium *E. coli*, the system as constructed may have broad utility across species. Our path to optimizing Mobile-CRISPRi for *Z. mobilis* revealed valuable lessons that may be generalizable across species: first, identify limiting components (either dCas9 or sgRNAs) by overexpression, and second, design strong synthetic promoters that take advantage of conserved interactions between RNA polymerase holoenzyme and DNA (47, 48) to improve portability.

CRISPRi is the ideal genetic tool to take advantage of the unusual properties of *Z. mobilis*. Because many metabolic genes are predicted to be essential, controlling metabolic flux may require constructing strains with partial knockdowns of essential genes. Mismatch-CRISPRi libraries are particularly well suited for empirically defining relationships between knockdown of metabolic genes (and associated changes in flux) with fitness, as strains comprising knockdown gradients of metabolic genes can be pooled and tested under a variety of growth conditions with fitness measured by next generation sequencing of sgRNA spacers. Further, the *Z. mobilis* chromosome is possibly polyploid (28, 29), or at least capable of duplicating at high frequency; this can cause problems with deletion/transposon insertion analysis of essential genes or other genes that have a strong impact on fitness. For instance, a high-throughput analysis of isolated transposon insertion mutants revealed that there was no significant difference in the probability of a transposon inserting a predicted essential versus non-essential gene (29), suggesting a polyploid chromosome and underscoring issues with interpreting insertion/deletion results in *Z. mobilis*. In contrast, CRISPRi is largely unaffected by polypoidy—it is capable of targeting essential genes across multiple copies of the chromosome as long as the sgRNA-dCas9 complex is expressed at high enough levels to account for multiple targets.

Numerous studies have linked hopanoid production and ethanol tolerance in *Z. mobilis* (27, 30, 56), but whether hopanoids provide resistance to non-physiological alcohols, such as IBA, remains unclear. Our CRISPRi results suggest that wild-type levels of extended hopanoids do not impart IBA resistance, but instead that preventing synthesis of extended (C_35_) hopanoids by blocking *hpnH* expression causes sensitivity. The simplest explanation for these results is that core C_30_ hopanoids, diploptene and diplopterol, accumulate in the cell and negatively impact the outer membrane. Further evaluation of this hypothesis will require careful tracking of the relationships between knockdown extent, fitness, IBA concentration, and levels of individual hopanoid species.

Our optimized CRISPRi system opens the door to high-throughput, systematic analysis of gene function in *Z. mobilis*. We envision that such screens will be invaluable for identifying genes involved in resistance to hydrolysate or biofuel inhibitors, genetic fingerprinting of hydrolysates from different plant sources or environments, and improving our understanding of the unique metabolism of *Z. mobilis*. We anticipate that this information will power the next generation of biofuel production strains, resulting in higher yields of advanced biofuels and bioproducts.

## MATERIALS AND METHODS

### Strains and growth conditions

Strains are listed in Table 1. *Escherichia coli* was grown in LB broth, Lennox (BD240230; 10 g tryptone, 5 g yeast extract, 5 g NaCl per liter) at 37°C aerobically in a flask with shaking at 250 rpm, in a culture tube on a roller drum, or in a deep 96 well plate with shaking at 200 rpm. *Zymomonas mobilis* was grown in DSMZ medium 10 (10 g peptone and 10 g yeast extract per liter plus 2% glucose) at 30°C aerobically without shaking. Media was solidified with 1.5% agar for growth on plates. Antibiotics were added when necessary: *E. coli* (100 μg/ml ampicillin, 20 μg/ml chloramphenicol, 50 μg/ml kanamycin) and *Z. mobilis* (100 μg/ml chloramphenicol, 120 μg/ml kanamycin). Diaminopimelic acid (DAP) was added at 300 μM to support growth of *dap*-*E. coli* strains. 0.1-1 mM Isopropyl β-d-1-thiogalactopyranoside (IPTG) was added where indicated. All strains were preserved in 15% glycerol at −80°C.

### Plasmid construction

Plasmids and construction details are listed in Table 2 and a representative plasmid map is shown in Fig. S7. *pir*-dependent plasmids were propagated in *E. coli* strain BW25141 (sJMP146) and other plasmids in *E. coli* strain DH10B (sJMP006). Plasmids were assembled from fragments (linearized vector, PCR products, and/or synthetic DNA) using the NEBuilder Hifi DNA assembly kit (New England Biolabs (NEB) E2621). Plasmids were cut with restriction enzymes from NEB. Linearized plasmids were re-ligated using T4 DNA ligase (NEB M0202). Fragments were amplified using Q5 DNA polymerase (NEB 0491) followed by digestion with DpnI. Fragments were purified using the Monarch PCR & DNA Cleanup Kit (NEB T1030) after digestion or amplification. Plasmids were transformed into electrocompetent *E. coli* cells using a BioRad Gene Pulser Xcell on the EC1 setting. Plasmids were purified using the GeneJet Plasmid Miniprep kit (Thermo K0503) or the Purelink HiPure Plasmid Midiprep kit (Invitrogen K210005). Site-directed mutagenesis of plasmids was performed by DNA synthesis with 2.5 U PfuUltra II Fusion HS DNA Polymerase (Agilent), 0.2 μM oligonucleotide encoding the change, 0.2 mM dNTPs, and 50 ng of plasmid DNA in a 25-μL reaction with a 1-min/kb extension time at 68°C, followed by DpnI digestion. sgRNA-encoding sequences were cloned into CRISPRi plasmids between the BsaI sites with inserts prepared by one of two methods. In method one, two 24 nt oligonucleotides were designed to overlap such that when annealed, they have ends that are complementary to the BsaI-cut ends on the vector. Oligos (2 μM each) were annealed in 1X CutSmart buffer (NEB) at 95°C for 5 min followed by cooling to room temperature. For method two, fragments were amplified by PCR with primers oJMP197 and oJMP198 from a 78 nt oligonucleotide followed by digestion with BsaI-HF-v2 and purification with the Monarch DNA purification kit (NEB) following the manufacturer’s oligonucleotide purification protocol. Inserts (2 μl of a 1:40 dilution of annealed oligos or 2 ng purified digested PCR product) were ligated into 50 ng BsaI-digested vector. Oligonucleotides and synthetic DNA gBlocks were purchased from Integrated DNA Technologies (Coralville, IA). Sequencing was performed by Functional Biosciences (Madison, WI).

**Table 2.**
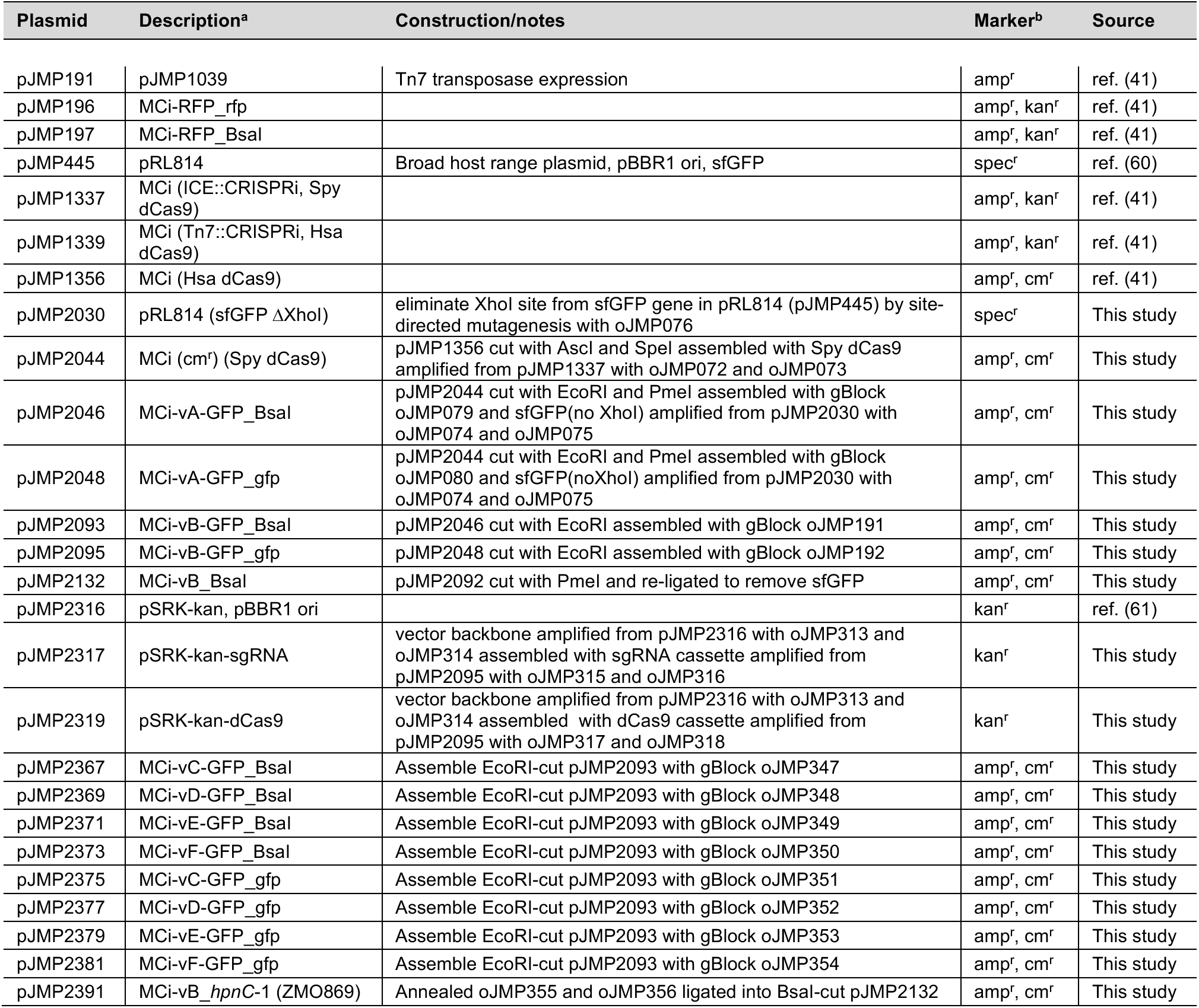
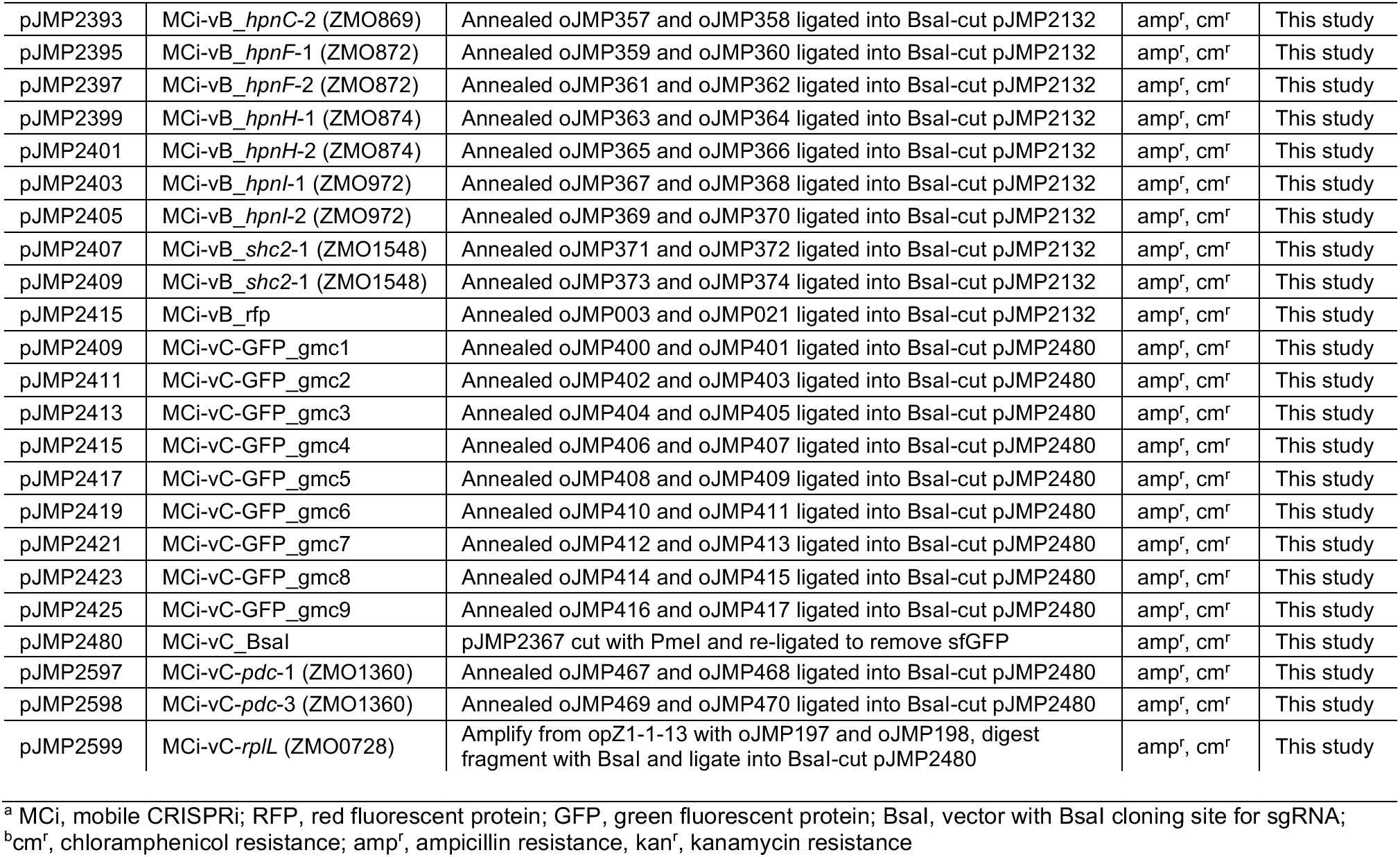
Plasmids.

### Transfer of CRISPRi system to *E. coli* and *Z. mobilis*

Strains with a chromosomally-located CRISPRi expression cassette were constructed by tri-parental mating of two donor strains – one with a plasmid encoding Tn7 transposase and another with a plasmid containing a Tn7 transposon encoding the CRISPRi system – and a recipient strain (either *E. coli* BW25113 or *Z. mobilis* ZM4 (PK15436)). The method was as described in (Peters 2019) with several modifications. Briefly, all matings used *E. coli* WM6026 which is *pir*^+^ to support *pir*-dependent plasmid replication, *dap*^−^ making it dependent on diaminopimelic acid (DAP) for growth and encodes the RP4 transfer machinery required for conjugation. Donor strains were grown ~16 h at 37°C in LB + 100 μg/ml ampicillin and 300 μM DAP. *E. coli* recipient was grown ~16 h at 37°C in LB. *Z. mobilis* recipient was grown ~24-30 h at 30°C in DSMZ10. Cells were centrifuged at 4000 × g for 5 min and gently resuspended twice in an equal volume of fresh medium with no antibiotic or DAP. For *E. coli* recipients, 700 μl LB, and 100 μl each donor and recipient was mixed in a sterile 1.5 ml microfuge tube. For *Z. mobilis* recipients, 200 μl DSMZ10, 500 μl *Z. mobilis*, 200μl transposase donor and 100 μl transposon donor was mixed in a sterile 1.5 ml microfuge tube. Cells were centrifuged at 4000 × g for 3 min, gently resuspended in 25 μl LB or DSMZ10, and pipetted onto a 13 mm cellulose filter placed on prewarmed agar plate (LB for *E. coli* or DSMZ10 for *Z. mobilis*). Plates were incubated at 37°C for 2-6 h for *E. coli* and 30°C for 24 h for *Z. mobilis.* After the incubation period, using sterile forceps, filters were placed into sterile 1.5 ml microfuge tubes containing 200 μl sterile 1X PBS, vortexed 20 s to dislodge cells from filters, diluted in 1X PBS and plated on appropriate medium for recipient and antibiotic to select for transposon (see above) with no DAP (to select against the donor). Efficiency of transposition was generally ~1 in 10^3^ for *E. coli* and ~1 in 10^5^-10^6^ for *Z. mobilis*. Isolated colonies were generally obtained from ~10-100 μl of 1:100 dilution per plate and isolated colonies were restruck for isolation to ensure purity.

### CRISPRi insertion onto *Z. mobilis* chromosome

Insertion of the CRISPRi expression cassette into the Tn7att site downstream of *glmS* in *Z. mobilis* was confirmed by PCR with primers oJMP057 and oJMP058 (flanking insertion site) and oJMP059 and oJMP060 (upstream of insertion site and within CRISPRi transposon).

### CRISPRi stability in *Z. mobilis*

Z. *mobilis* strains with chromosomally located CRISPRi expression cassettes (6 individual isolates) were grown in liquid culture medium with antibiotic selection to saturation. This culture was serially diluted 10-5 into non-selective medium (starting OD A_600_ ~0.00002) and grown ~17 generations back to saturation (OD A_600_ ~2.0). Dilution and growth was repeated 2 additional times for a total of ~50 generations prior to plating on non-selective plates. Forty-eight isolated colonies were selected and patched on selective and non-selective plates and all strains retained the ability to grow on the antibiotic whose resistance was conferred by the chromosomally located CRISPRi expression cassette.

### GFP/RFP knockdown assays

GFP or RFP knockdown was measured using a plate reader (Tecan Infinite 200 Pro M Plex). Cell density was determined by absorbance at 600 nm (A_600_) and fluorescence was measured by excitation/emission at 482/515 nm for GFP and 555/584 nm for RFP. Initial cultures (n=4) were grown from single colonies to saturation (~30 h for *Z. mobilis* or ~16 h for *E. coli*) in 1 ml medium in 96 well deepwell plates. These cultures were serially diluted 1:1000 (*Z. mobilis*) or 1:10,000 (*E. coli*) into 1 ml fresh medium (no antibiotic and 0-1 mM IPTG as indicated) and grown back to saturation (~24-30 h for *Z. mobilis* or ~8-16 h for *E. coli*). Pelleted cells were resuspended in 1 ml 1X PBS, diluted if necessary, and 200 μl was transferred to a clear bottom black microtiter plate and measured in the plate reader as indicated above. Fluorescence values were normalized to cell density and to measurements from strains not expressing GFP.

### Gene knockdown spot dilution assay

Z. *mobilis* strains with chromosomally located CRISPRi expression cassettes (2 individual isolates) were grown in liquid culture medium with antibiotic selection to saturation. These cultures were serially diluted 1:10 in non-selective medium and 3 μl was spotted onto plates containing 0, 0.1 mM or 1 mM IPTG which were incubated at 30°C aerobically prior to analysis.

### Gene knockdown growth assay

Z. *mobilis* strains with chromosomally located CRISPRi expression cassettes (n=2 individual isolates) were grown in liquid culture medium with antibiotic selection to saturation. These cultures were serially diluted 1:1000 into non-selective medium with 0 or 0.1 mM IPTG and 0, 0.63%, 1.25% or 2.5% isobutanol in a 96 well deep well plate and incubated at 30°C aerobically prior to analysis of growth (measured as absorbance at 600 nm (A_600_)).

**Table 3.**
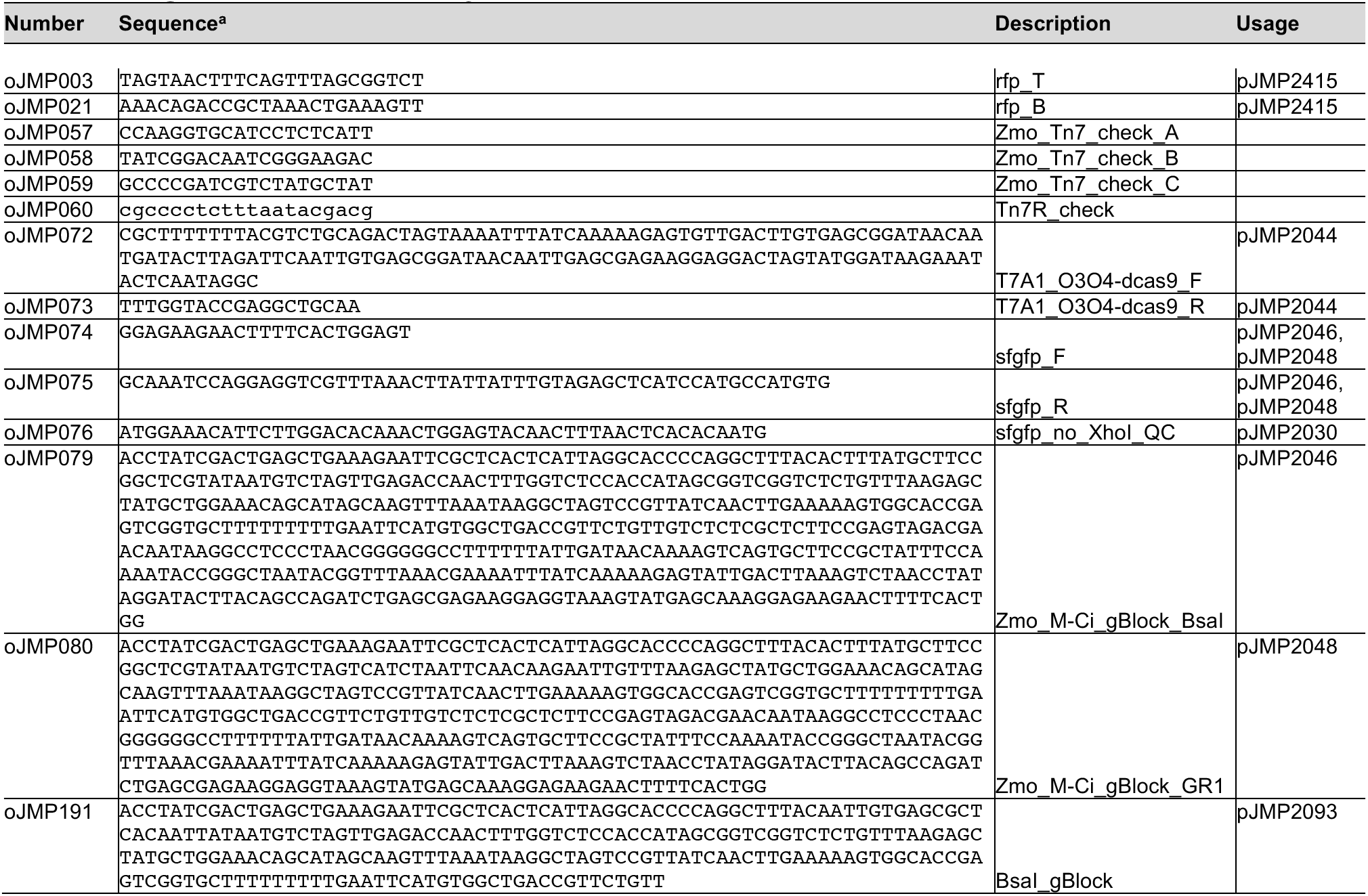
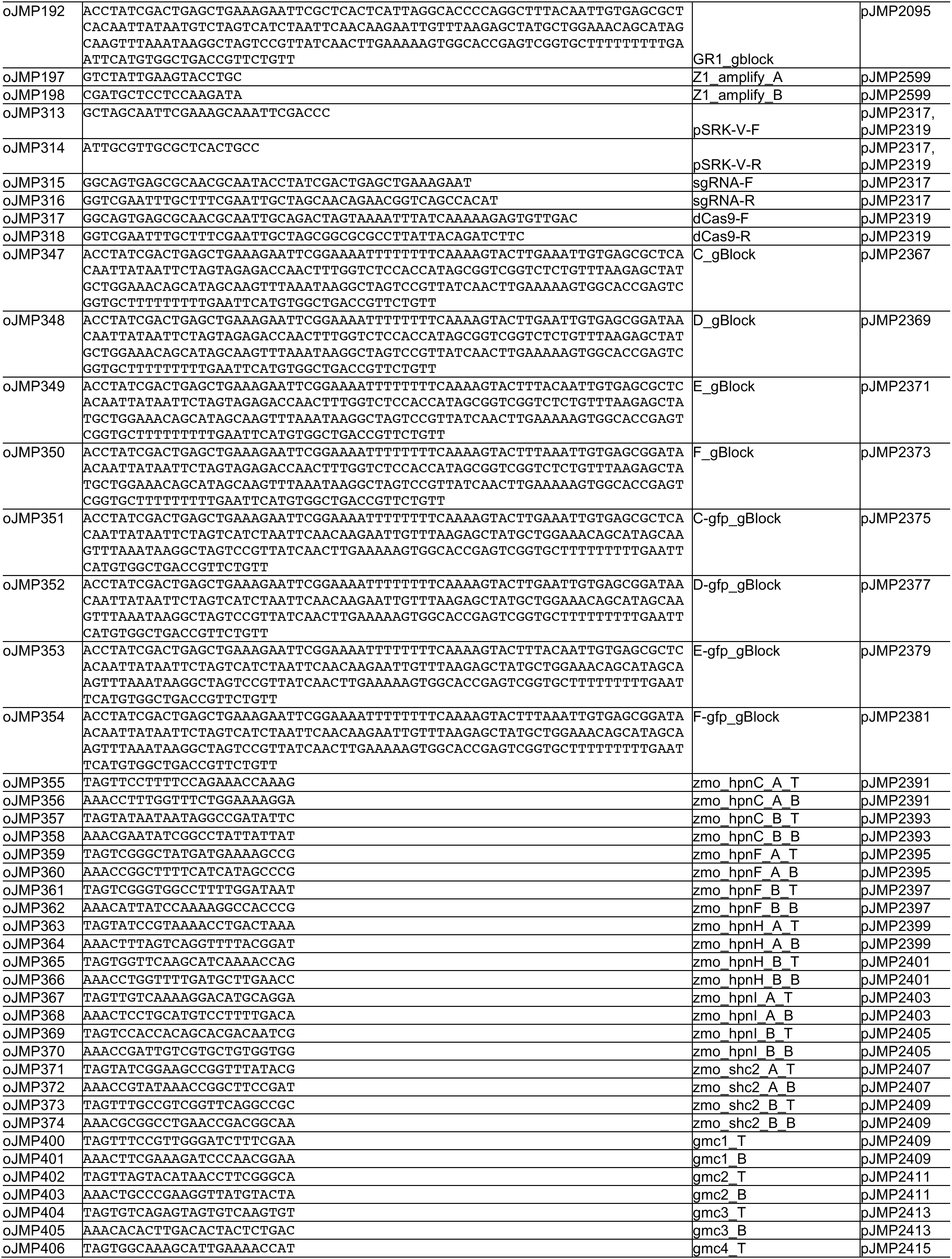
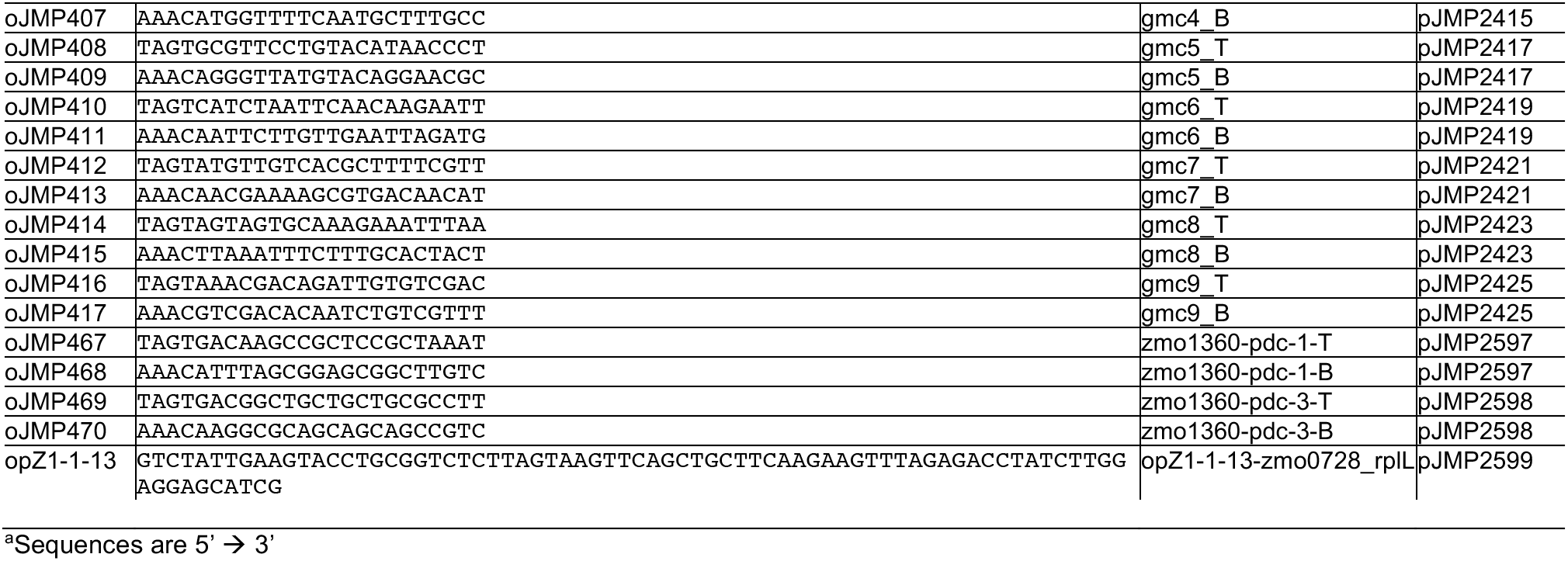
Oligonucleotides and Synthetic DNA.

## Data Availability

Plasmids and their sequences are available from Addgene under accession numbers xxxx-xxxx (note: accession #s pending).

## ACKNOWLEDGEMENTS

We thank members of the TerAvest, Landick, Kiley, Amador-Noguez, Reed, Sato, Hittinger and Gross labs for helpful feedback. We thank Yang Liu, Robert Landick, Piyush Lal, and Tricia Kiley for strains and plasmids. This work was supported by the U.S. Department of Energy, Office of Science, Office of Biological and Environmental Research, Great Lakes Bioenergy Research Center under Award Numbers DE-SC0018409 and DE-FC02-07ER64494.

---

**Fig S1.**
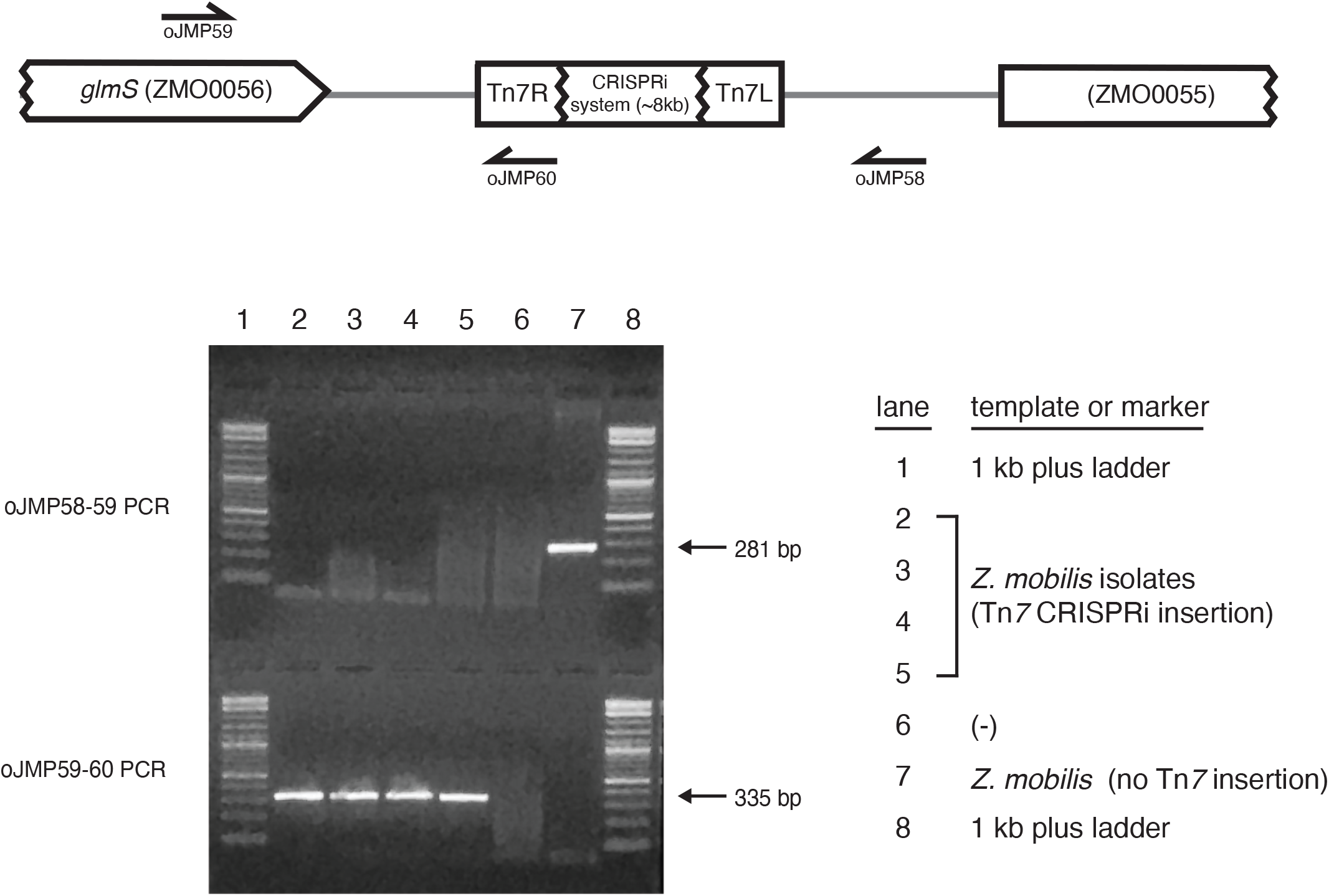
PCR test to confirm correct insertion of Mobile-CRISPRi into the *Z. mobilis* genome downstream of *glmS*.

**Fig S2.**
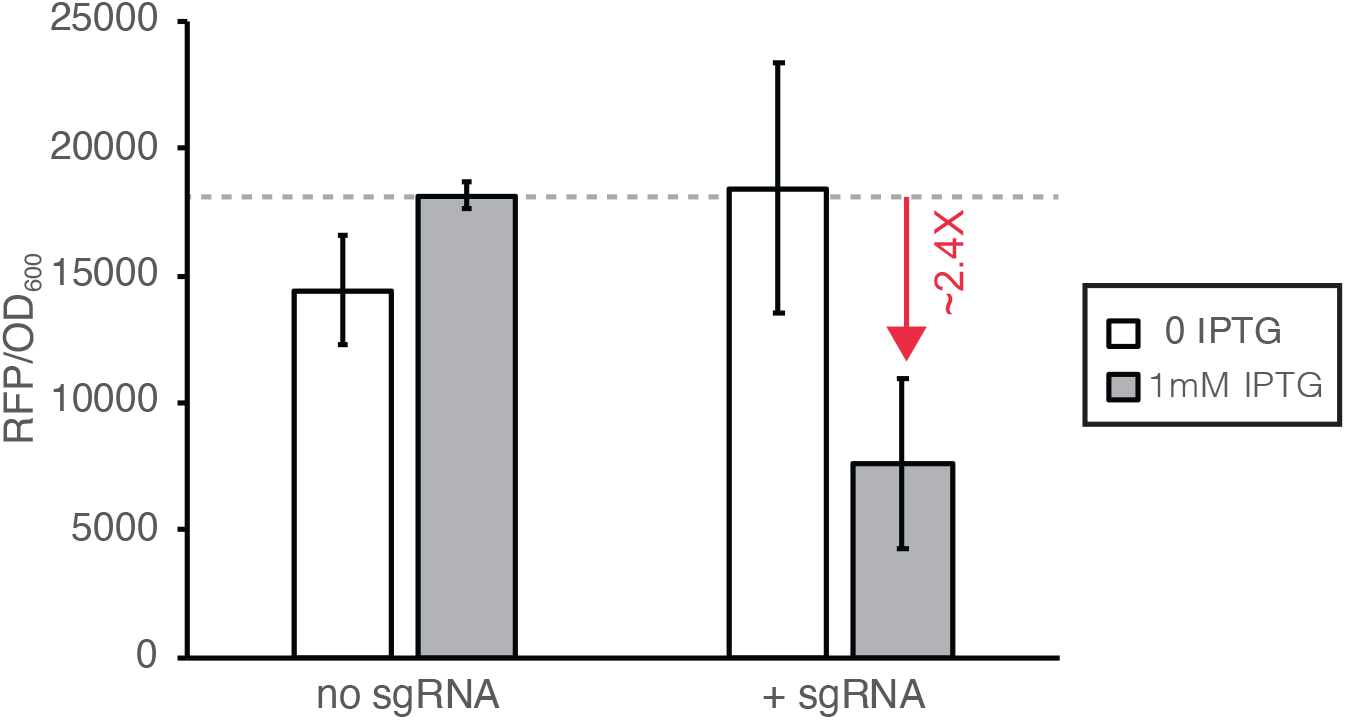
Knockdown of RFP using the Mobile CRISPRi system from Peters, et al. 2019 prior to optimization for *Z. mobilis*. Red arrow indicates fold change compared to control. Growth was measured by absorbance at 600nm.

**Fig S3.**
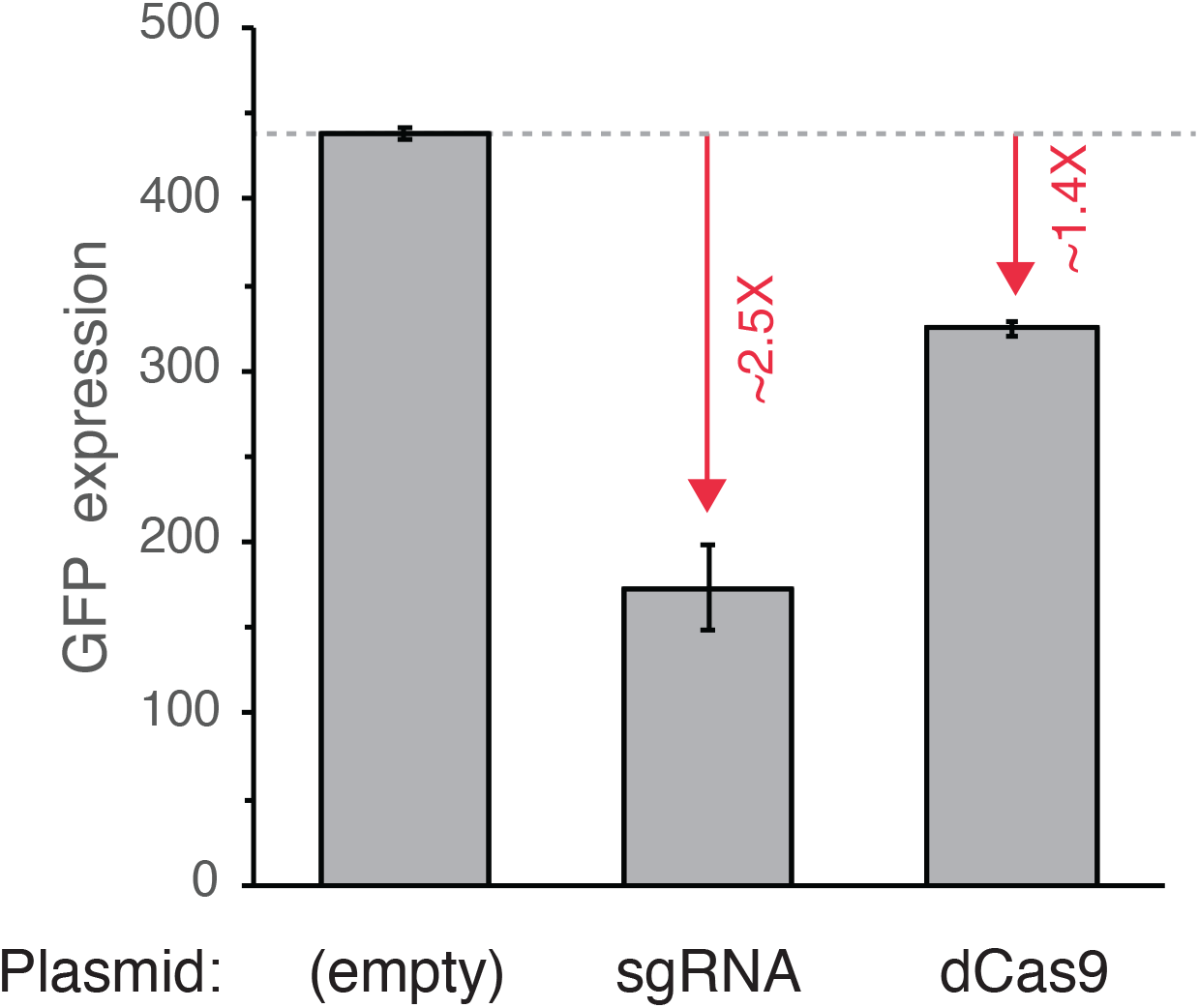
sgRNA is limiting in *Z. mobilis* promoter variant B CRISPRi system. Knockdown of GFP in *Z. mobilis* with the promoter variant B Mobile CRISPRi system on the chromosome and either additional sgRNA or additional dCas9 expressed from the broad host range plasmid pSRK-kan. Red arrows indicate fold change compared to control.

**Fig S4.**
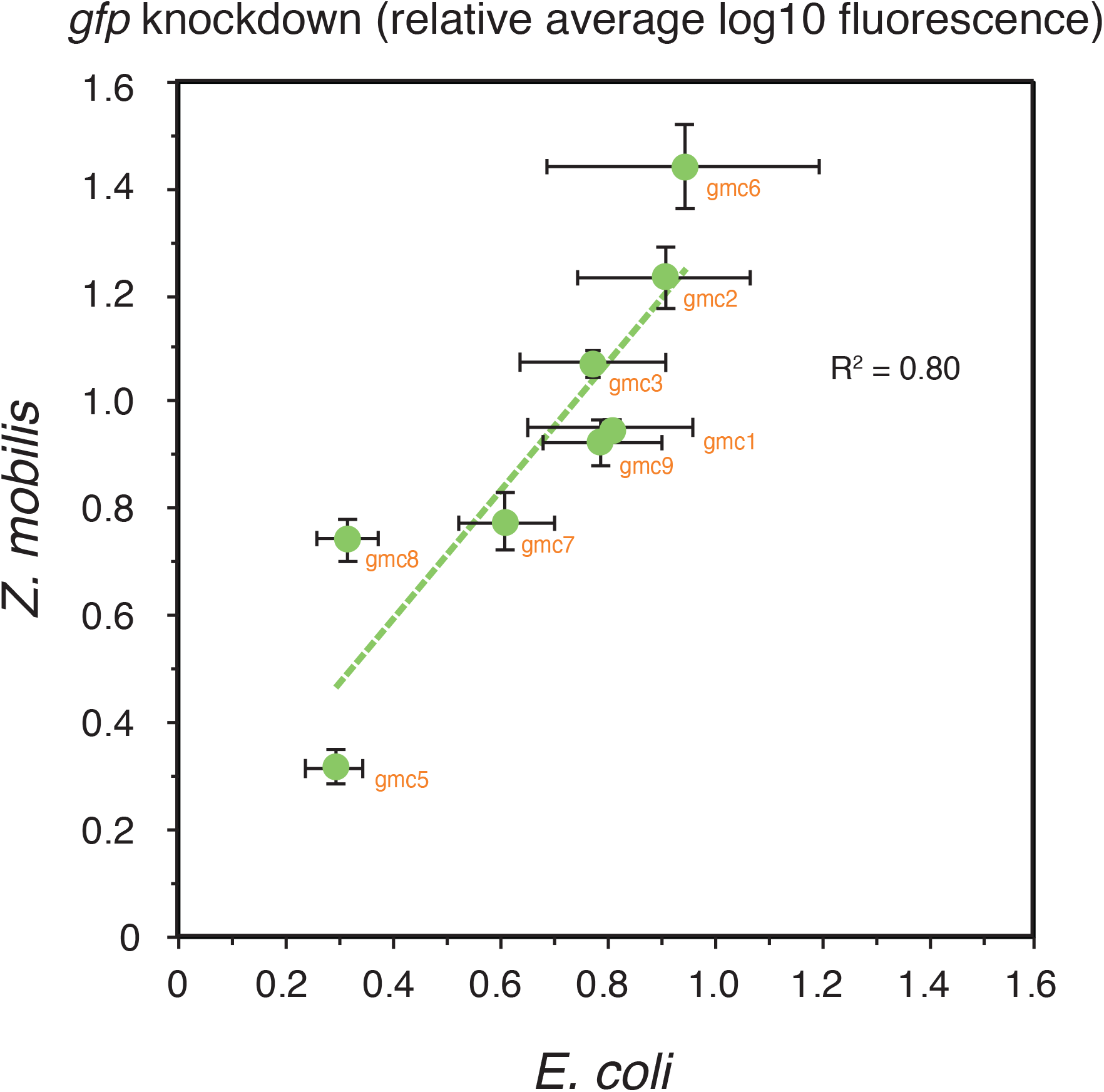
Comparison of *gfp* knockdown by mismatch sgRNAs in *Z. mobilis* vs *E. coli*. Graph shows *gfp* knockdown (relative average log10 fluorescence) of *Z. mobilis* vs E. coli CRISPRi strains expressing sgRNAs targeting *gfp* (indicated in orange). Linear trendline is dashed green.

**Fig S5.**
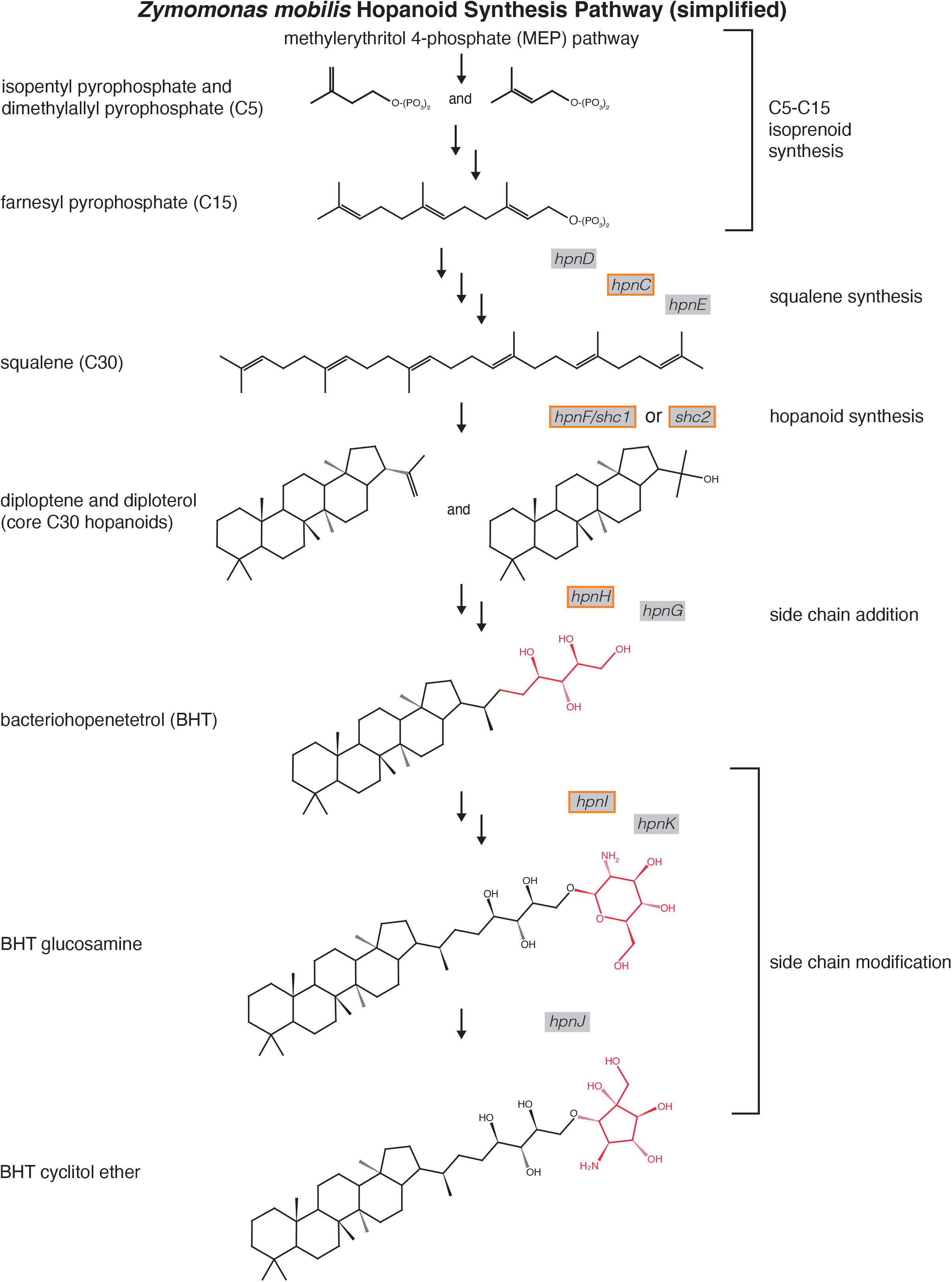
Hopanoid lipid synthesis pathway in *Z. mobilis*. Relevant lipids and their structures are shown (some intermediates omitted for simplicity). Genes encoding enzymes are indicated in grey boxes next to the arrows, with CRISPRi target genes boxed in orange. Red is used to highlight the change from the previous structure.

**Fig S6.**
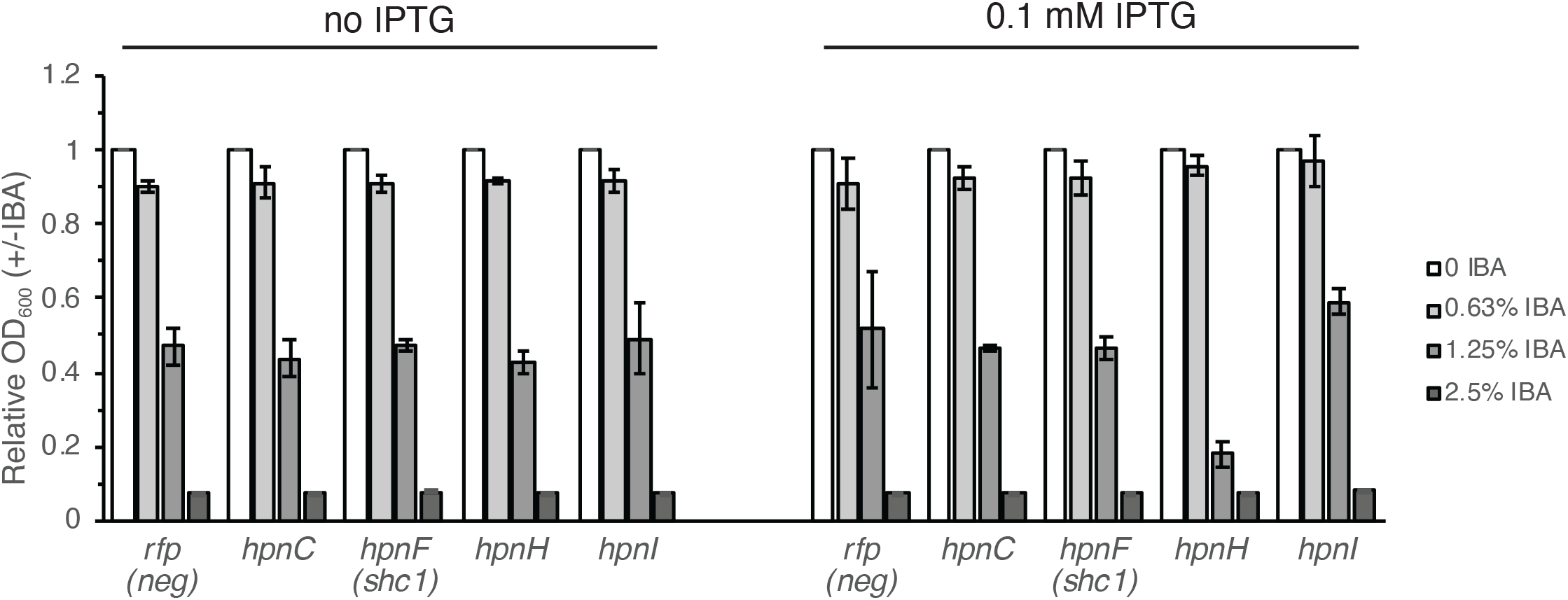
CRISPRi knockdown of hopanoid lipid synthesis related genes. Expanded version of data in Fig 4C showing additional concentrations of isobutanol (0.63%, 1.25%, 2.5% and 0 or 0.1 mM IPTG. Standard deviation between 4 replicates is shown.

**Fig S7.**
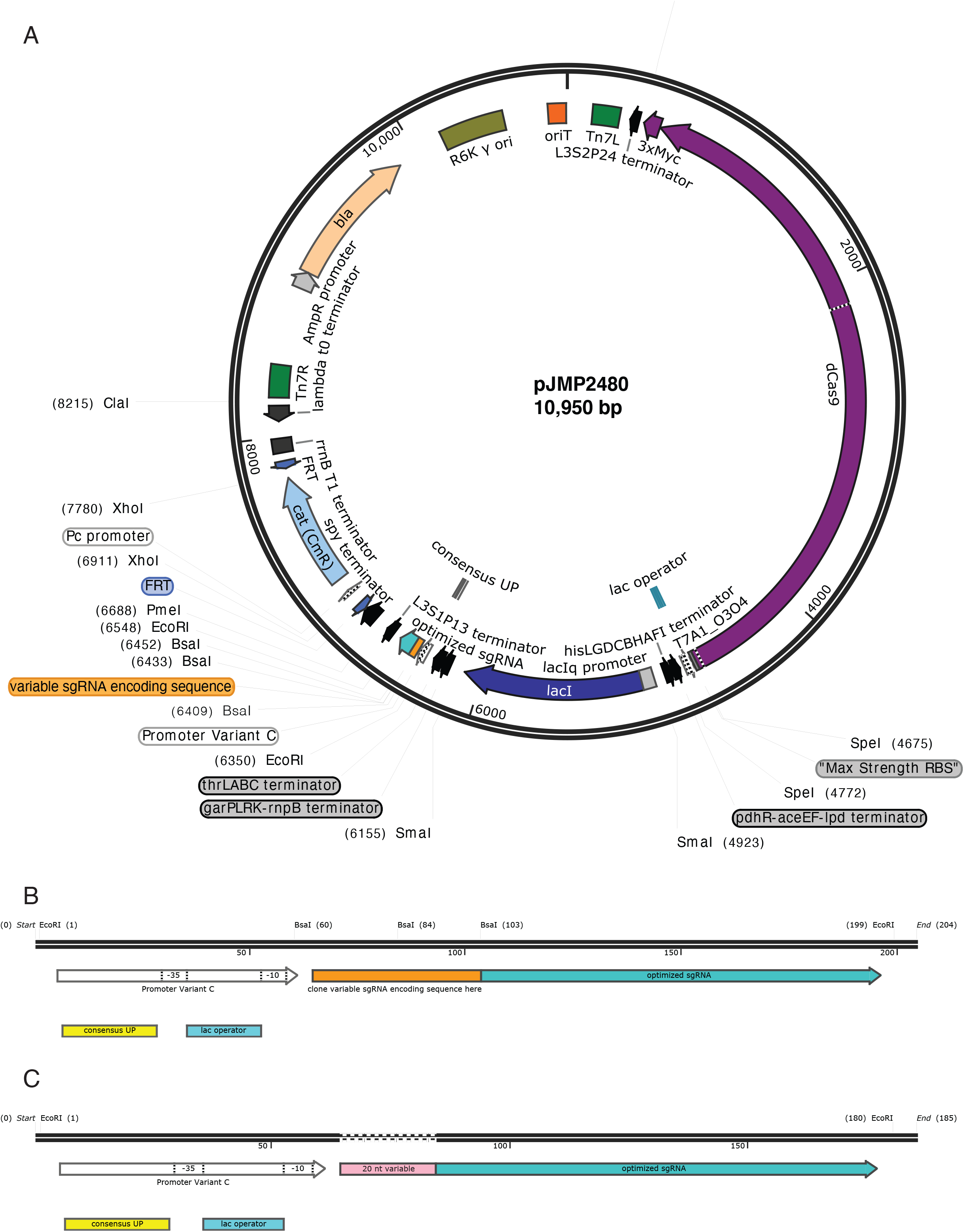
Mobile CRISPRi system for *Zymomonas mobilis*. (A) Plasmid map of pJMP2480 showing CRISPRi system located on a transposon (between Tn7L and Tn7R in dark green) that can be transferred to *Z. mobilis* genome via Tn7 transposase-mediated transposition. The modular design enables parts to be exchanged as necessary by restriction enzyme digest (restriction enzyme sites shown) followed by Gibson assembly. All components are separated by transcriptional terminators. Once integrated into the genome, the system is stable without antibiotic selection for > 50 generations. The *cat* antibiotic cassette (light blue) is flanked by FRT sites (dark blue) to enable removal from the genome using Flp-recombinase, if desired. Full sequence of the plasmid is available at addgene.org accession # xxxx. (B) Zoomed in view of the sgRNA expression cassette (between EcoRI sites). The 20 nt variable region of the sgRNA can be cloned between the BsaI sites. (C) Zoomed in view of the sgRNA expression cassette showing location of the 20 nt variable region (pink).

